# A neuroprotective locus modulates ischemic stroke infarction independent of collateral vessel anatomy

**DOI:** 10.1101/2020.12.04.411397

**Authors:** Han Kyu Lee, Sarah E. Wetzel-Strong, David L. Aylor, Douglas A. Marchuk

**Affiliations:** Department of Molecular Genetics and Microbiology, Duke University Medical Center, Durham, North Carolina, United States of America; Department of Biological Sciences, North Carolina State University, Raleigh, North Carolina, United States of America

**Author notes:** Corresponding author (DAM); (HKL).

## Abstract

Ischemic stroke is caused by a disruption of the blood supply to the brain leading to neuronal cell death. Genetic studies of ischemic stroke have identified numerous gene variants that increase the risk to develop stroke. In stark contrast, genetic studies of stroke outcomes, such as the infarct territory size, are confounded by many uncontrollable variables, leading to a paucity of gene targets for treatment of an incipient stroke. Using genetically diverse inbred strains of mice and a surgically-induced model of ischemic stroke, we used quantitative trait locus mapping to identify novel gene targets modulating infarct size, which varies greatly across inbred strains. Although infarct size is largely determined by the extent of collateral vessel connection between arteries in the brain that enables reperfusion of the ischemic territory, we have identified strain pairs that do not vary in this phenotype, but which nonetheless exhibit large differences in infarct size. In this study we performed QTL mapping in mice from an intercross between two such strains, WSB/EiJ and C57BL/6J. We identified a strong locus on chromosome 8 that overlaps with a locus of similar direction and effect previously mapped in an intercross between C3H/HeJ and C57BL/6J strains. To identify causative genes within the overlapping genetic interval, we surveyed nonsynonymous coding SNPs and performed RNA sequencing data analysis for all three mapping strains. We identified Macrophage Scavenger Receptor 1 (*Msr1*) as a strong candidate gene that harbors multiple coding SNPs predicted to be damaging. Using *Msr1*-deficient mice, we demonstrated that cerebral infarct volume after stroke induction is dramatically increased in a strain background where reperfusion effects due to collateral vessels is blunted. Significantly, the identification of neuroprotective genes such as *Msr1* provides new genes for future mechanistic studies of infarction following ischemic stroke and provides novel gene/protein targets for therapeutic development.

**Author summary:** The most common form of stroke arises when a blockage occurs in a blood vessel of the brain, thereby preventing delivery of oxygen and nutrients to areas supplied by the affected vessel, leading to tissue death. The main treatment for this form of stroke is medication to dissolve the blockage; however, more treatment options are required to better reduce the death and disability associated with stroke. In this study, we sought to identify genes that can decrease the amount of damage to brain tissue following a stroke, with a specific focus on examining genes that work to directly protect the neurons, rather than returning blood flow to the affected area. Since it is impossible to precisely control the nature of stroke and the genetic variability in humans, we used mice identify a genetic region that is associated with the amount of tissue damage following stroke. Within this genetic region, we identified a list of candidate genes, including the gene *Msr1*, which we found is important for controlling tissue damage in one genetic population of mice after stroke. The genes identified here require further follow-up to determine the impact on stroke outcomes and the usefulness of these candidates as therapeutic targets.

## Introduction

Ischemic stroke is caused by a disruption of blood supply to the brain leading to neuronal cell death in the ischemic region. Fifteen million people worldwide suffer a stroke [1-3], with the wordwide health care costs estimated at $65.5 billion [4]. In the US, stroke is the fourth-leading cause of death with almost 800,000 new or recurrent cases occurring each year [5]. Risk factors for stroke include environmental influences (e.g., smoking, diabetes, obesity, etc.), but epidemiologic studies have estimated that two-thirds of the risk to develop ischemic stroke is due to genetic factors [6,7]. GWAS studies for ischemic stroke, including a recent meta analysis, have identified dozens of variants in genes involved in diverse biological processes, and many variants share risk factors with other vascular traits or intermediate phenotypes of stroke [8]. Pharmacological intervention to reduce stroke *risk* is focused on reduction of these related vascular risk factors [8].

By contrast, once a stroke has occurred, the only approved drug for stroke therapy is intravenous recombinant tissue plasminogen activator (tPA), currently given to only 2-3% of stroke patients because of its limited time window for administration and major risk for adverse effect. Unfortunately, tPA does not provide protection to neural tissues already damaged by stroke. Thus, there is an urgent need to identify and develop new drug targets to provide effective neuroprotection and/or neuroresuscitation to damaged tissues. Identifying such gene targets has been problematic. In contrast to the successes of genetic studies of stroke *risk*, GWAS of stroke outcomes among ischemic stroke patients are intrinsically difficult due to uncontrollable variation in the extent and location of the occluded vessel, strong environmental variables (e.g., diet, smoking, etc.), and especially variation in the critical time between first recognized symptoms of stroke and medical intervention.

We have attempted to identify genes that influence the critical ischemic stroke outcome, infarct volume, using a surgically-induced mouse model of ischemic stroke and QTL mapping analysis. We have identified several genetic loci that regulate infarct volume [9-11] and identified several genes modulating this trait [10,12,13]. The strongest locus controlling infarct volume in mice [10] also modulates the collateral vascular anatomy [14], and a gene underling both phenotypes was shown to be *Rabep2* [15]. The collateral circulation enables vascular reperfusion of the ischemic territory, thus diminishing the effects of ischemia. However, as the collateral vessel anatomy is established early in life due to genetic factors [11,16], we sought to identify genes that modulate infarct volume, independent of the collateral circulation in the hopes of identifying loci and genes that modulate infarct size through an innate, neuroprotective effect. Our strategy is to perform QTL mapping in crosses between inbred mouse strains that exhibit similar collateral vessel anatomy, but which nonetheless show large differences in infarct volume after ischemic stroke. In our survey of strain phenotypes we have included the eight founder mouse strains of the Collaborative Cross (CC) that include three wild-derived strains, CAST/EiJ (CAST), PWK/PhJ (PWK), and WSB/EiJ (WSB), containing extensive diversity compared to classical inbred mouse strains [17-19]. Candidate genes mapping within linkage peaks are identified through ancestral haplotype analysis (when crosses with multiple inbred strains uncover the same locus), strain-specific RNA expression analysis, and *in silico* analysis of coding variants. Highly compelling candidate genes are then validated with phenotypic studies using genetically modified mice. Through this approach, we have identified loci and genes that modulate infarct volume independent of the effects of the collateral vasculature [10-13,20].

In the present study, by performing QTL mapping between a wild-derived inbred strain, WSB, and common classical inbred strain, C57BL/6J (B6), we identify a neuroprotective locus that overlaps with a locus mapped in an independent QTL analysis between two classical inbred strains, C3H and B6. Using reciprocal congenic mouse lines, we validate this locus modulating infarct volume via a collateral-independent manner. By examining non-synonymous coding SNP variation and strain-specific differential gene expression, we identify potential candidate genes within the locus. We further investigate one of these genes, *Msr1*, using a knockout allele, and demonstrate that this gene product modulates infarct volume in a collateral-independent manner.

## Results

### A locus on chromosome 8 regulates infarct volume in a cross between two inbred mouse strains, B6 and WSB

To map additional genetic loci regulating infarct volume, focusing on strains that would be likely to exhibit differences due to vascular independent effects, we returned to our phenotypic characterization of the eight founder mouse strains of the CC [20]. As predicted by the role of collateral vessels in the reperfusion of ischemic tissue, classical inbred strain B6 exhibits a high number of collateral vessel connections with a correspondingly small infarct volume. By contrast, although a wild-derived strain WSB shows an even greater number of collateral vessel connections than B6, WSB mice exhibit a much larger infarct volume. This break between the collateral vessel density and infarct volume phenotypes in WSB motivated the choice of WSB and B6 for a new mapping cross.

To discover the genetic region(s) modulating ischemic infarction across these strains, we first generated F1 progeny between the strains, and then performed an intercross. Then, F1 and F2 animals were analyzed for collateral vessel and infarct volume phenotypes. The number of collateral vessel connections in the F1 animals was tightly distributed approximately midway between the parental strains (B6 (20.4), WSB (27.3), F1 (24.9)) (Fig 1A). Infarct volume in the F1 animals was also relatively tightly distributed, but much closer to the low values seen in B6 (B6 (7.8 mm^3^), WSB (22.6 mm^3^), F1 (8.9 mm^3^)) (Fig 1B). Collateral vessel number measured in a separate cohort of F2 animals was tightly distributed around the mean of 25.3. By contrast, infarct volume in the F2 animals exhibited a very wide distribution, ranging from 0.8 to 37.1 mm^3^ (Fig 1B). The tight distribution of collateral vessel numbers in the F2 animals, contrasted with the wide variation in infarct volume, validated our choice of these strains for our attempt to map collateral-independent loci that modulate infarct size.

**Fig 1.**
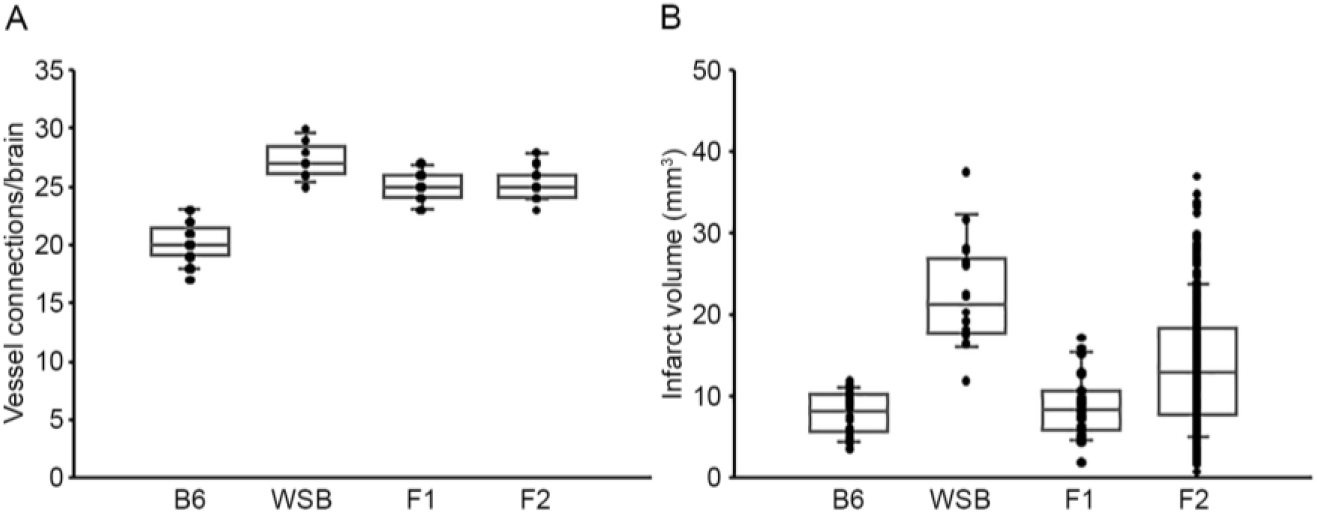
F2 progeny between B6 and WSB display a wide distribution of infarct volume after pMCAO. **(A)** The scatter plot indicates the number of collateral vessel connections between the ACA and MCA for parental strains, B6 and WSB, and their F1 and F2 intercross animals. The box plot shows the degree of dispersion and skewness. The number of animals for B6, WSB, F1, and F2 was 37, 13, 21, and 30 animals, respectively. Data represent the mean ± SEM. **(B)** The scatter and box plot graphs show the distribution of infarct volume after pMCAO for B6, WSB, and their F1 and F2 intercross animals. The number of animals for the infarct volume measurements was 34, 18, 27, and 376 animals, respectively. Data represent the mean ± SEM.

To map the genetic region(s) that modulate infarct volume in the cross, we performed genome-wide QTL mapping analysis. A total of 376 F2 mice (Fig 1B) were genotyped with a genome-wide mapping panel (the miniMUGA genotyping panel consisting of approximately 11K SNPs). For intitial genome-wide QTL mapping analysis, 515 informative SNP markers were selected across the mouse genome (S1 Table), which completely covered the entire mouse genome (markers were chosen at approximately 5 Mb intervals across each of the chromosomes). We identified a QTL peak on chromosome 8 (logarithm of the odds, LOD, 5.16; *p* = 0.002) that displayed significant linkage to the infarct volume trait (Fig 2). No other region reached statistical significance at *p* = 0.05, as determined by data permutation, but one peak on chromosome 17 reached a LOD of 3.51, corresponding to a suggestive *p* value of 0.083.

**Fig 2.**
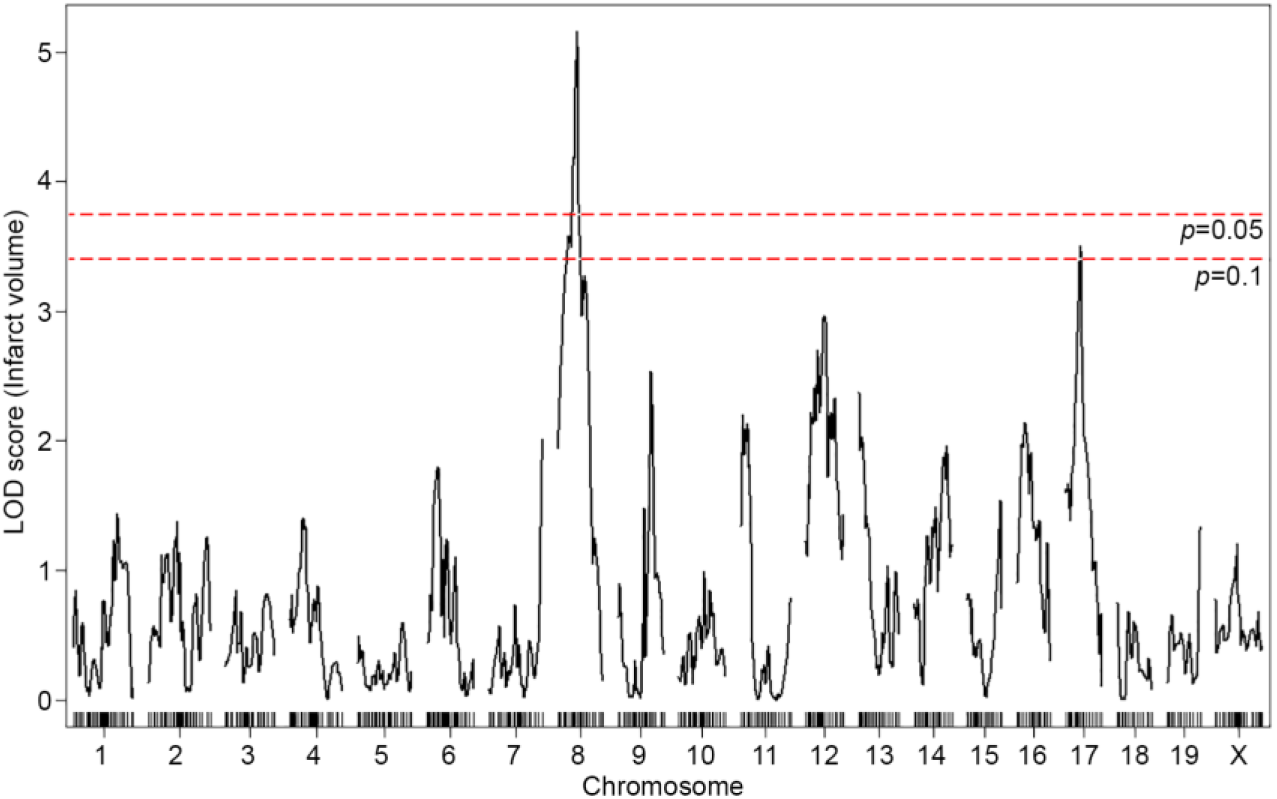
A locus for infarct volume is mapped in the F2 progeny between B6 and WSB. The graph presents the analysis of a genome-wide QTL mapping scan for infarct volume measured 24 hr after pMCAO using 376 F2 animals (B6 and WSB). Chromosomes 1 through X are represented numerically on the x-axis and the y-axis represents the LOD score. The significant (*p* < 0.05) level of linkage was determined by 1,000 permutation tests. Only a single genomic region on chromosome 8 displays significant linkage to infarct volume with a LOD score of 5.16.

The infarct volume phenotype of the F2 animals was relatively normally distributed, although with a tail at the high-end values (S1 Fig). To determine if deviations from normality were influencing our mapping analysis, we also performed non-parametric mapping. The linkage profiles across the entire genome were nearly identical, including at the linkage peaks on chromosomes 8 and 17 (S2 Fig). Thus, the mapping results appear robust to any deviations from normality.

### Characterization of the location, size and alleleic effects of the chromosome 8 locus

To fine-map the statistically-significant peak mapping to chromosome 8, we extended the interval an additional 1.5 Mb flanking each arm of the 1.5-LOD support interval; a convention that defines a conservative estimate of the locus. A total of 88 additional informative SNPs markers were included in the linkage data to more precisely map the locus. The resulting candidate interval falls between 36.16 and 78.11 Mb (Fig 3A and Table 1).

**Table 1.** Characteristics of two independent QTL mappings for surgically-induced ischemic infarct volume.

| Location | LOD score | 1.5-LOD interval (Mb) | Marker at peak<br>(position, Mb) | Mapping mouse strains | Protective<br>allele |
| --- | --- | --- | --- | --- | --- |
| Chr8 | 5.16 | 8: 36.16 - 78.11<br>(41.95 Mb) | gUNC14975151<br>(70.88 Mb) | B6 and WSB | WSB |
| Chr8 | 10.31 | 8: 36.02 - 50.26<br>(14.24 Mb) | rs13479735<br>(45.52 Mb) | B6 and C3H | C3H |

**Fig 3.**
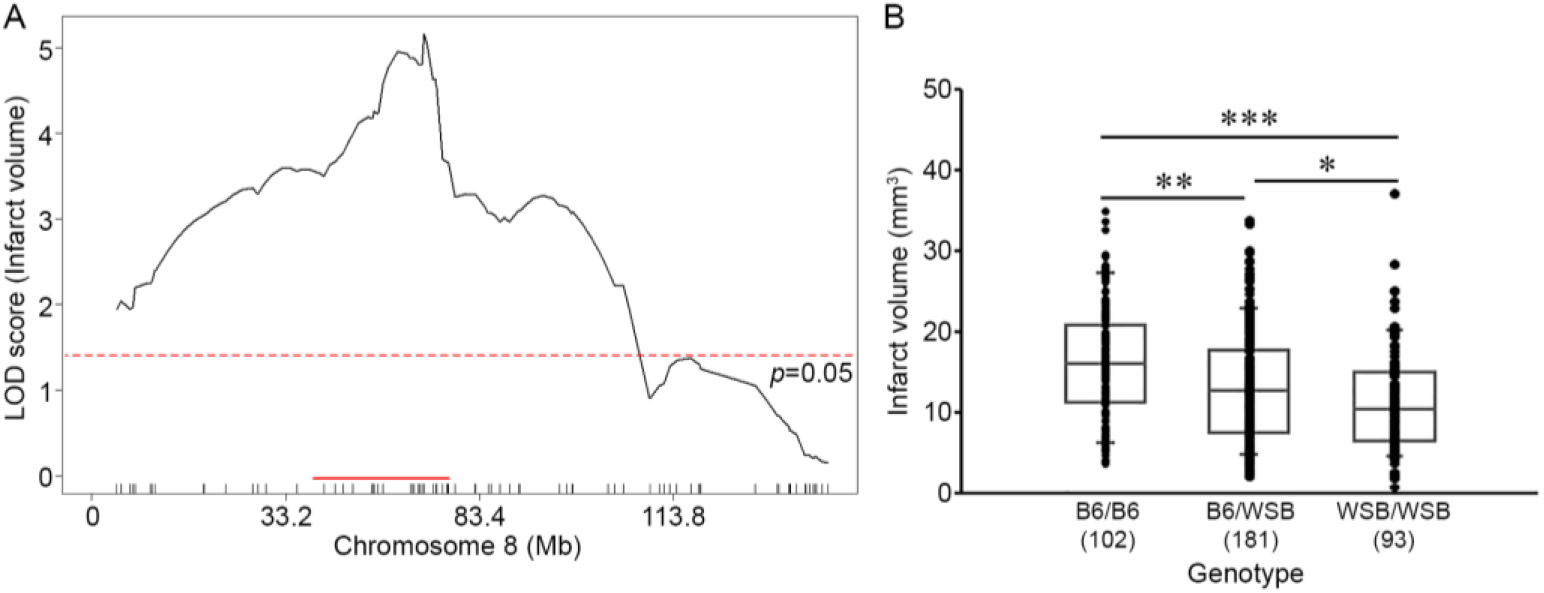
Characterization of the location, size, and allelic effects on infarct volume at the chromosome 8 locus. **(A)** The graph shows the QTL mapping across chromosome 8 using 88 informative SNP markers. The LOD score at the peak is 5.16 (gUNC14975151), and the 1.5-LOD support interval is from 36.16 to 78.11 Mb, indicated by the red bar on the graph. The significance threshold for linkage was calculated using the 88 informative SNP markers on chromosome 8. **(B)** Genotype-phenotype correlation of the F2 cohort at gUNC14975151. The B6 allele confers increased susceptibility to infarction and the WSB allele confers protection. Data represent the mean ± SEM. * *p* < 0.05 and *** *p* < 0.001, 2-tailed Student’s *t* test.

Surprisingly, the direction of the allele-specific phenotypic effect at this locus is opposite to that observed for the parental strains. Although infarct volume for B6 mice is smaller than that of WSB mice, in the F2 animals, B6 homozygotes at this locus exhibit larger infarct volumes than WSB homozygotes, with heterozygotes falling between these values (Fig 3B). Thus, unexpectedly, the WSB allele from the otherwise highly sensitive inbred strain confers resistance to ischemic tissue damage.

To survey how other loci affect infarct volume, we examined the allelic effects of the next highest peak, mapping to chromosome 17. Although this locus did not quite reach the significance threshold level (Fig 2), B6 homozygotes at this locus exhibit a protective effect on infarct volume, in agreement with the phenotype of the parental B6 strain (S3 Fig). These data suggest that effects of many different regions of the genome, including the chromosome 17 locus, confer the overall infarct volume phenotype of the WSB and B6 strains, and override the opposite allelic effects of a the locus mapping to chromosome 8.

### Concordance with a previously mapped QTL

The directional phenotypic effects at the chromosome 8 locus parallel those of another infarct volume QTL mapping to mouse chromosome 8. Like WSB, classical inbred mouse strain C3H appears to break the inverse correlation between collateral vessel density and ischemic damage, with a relatively high number of collateral vessel connections but showing a paradoxically large infarct volume. Infarct volume in the B6 strain (7.8 mm^3^) is approximately 3-fold smaller than that observed in both C3H (26.4 mm^3^) and WSB (22.6 mm^3^) strains (S4 Fig). In a cross between C3H and B6, we previously mapped a highly significant QTL to chromosome 8, where the direction of the allele-specific phenotypic effect at this locus was opposite to that observed for the parental strains [11]. This locus, named *Civq4*, overlaps the currently mapped locus between WSB and B6. Wild-derived strain WSB is more distantly related than either of the classical strains, providing a hypothesis that would aid in fine mapping the locus. If we assume the same QTL is segregating in both crosses, we would expect ancestral haployptes to follow a pattern of WSB = C3H ≠ B6 in the candidate region [21,22].

We thus re-analyzed our previous QTL mapping data between B6 and C3H to provide a robust comparison to the locus mapped in this study. Using 285 informative SNP markers along with the infarct volume phenotype of 210 F2 animals in the cross between B6 and C3H, we performed genome-wide QTL mapping analysis. Similar to the previous QTL mapping data [11], we observe a highly significant linkage peak (LOD 10.31) located on chromosome 8 (S5 Fig A). The 1.5-LOD support interval of *Civq4* (36.02 –50.26 Mb) falls completely within the 1.5-LOD support interval of this newly mapped locus (36.16 – 78.11 Mb) (Fig 3A, S5 Fig B, and Table 1). As previously noted [11] the allele-specific phenotypic effect of this locus is also opposite to that observed for the parental strains (S5 Fig C). Thus, in two independent QTL mapping experiments with three different inbred strains, we have identified a genetic region on chromosome 8 where the B6 allele confers increased sensitivity to infarction.

### Reciprocal congenic mouse lines validate the phenotypic effects of the QTL interval on chromosome 8

To validate the phenotypic effects of the previously-mapped *Civq4* locus, we generated reciprocal congenic mouse lines carrying a segment of the *Civq4* region either from the B6 introgressed into the C3H background (C.B6-*Civq4*; LineC) or from the C3H introgressed into the B6 background (C.C3H-*Civq4*; LineB). These lines were generated by repeated backcrosses (10 generations) selecting progeny at each backcross that retaining the *Civq4* locus from the relevant parental line while eliminating the rest of the genome from this parent. Congenic mouse LineB contains approximately 28.48 Mb of the C3H region of *Civq4* (from 31.34 to 59.82 Mb) in the B6 background and congenic mouse LineC contains approximately 23.83 Mb of the B6 region of *Civq4* (from 35.99 to 59.82 Mb) in the C3H background (Fig 4A). Operating under the assumption that the chromosome 8 locus mapped in the C3H x B6 cross [11] and the WSB x B6 cross are the same, the congenic lineB and LineC (from the C3H and B6 cross) further narrowed the candidate interval by approximately 50%.

**Fig 4.**
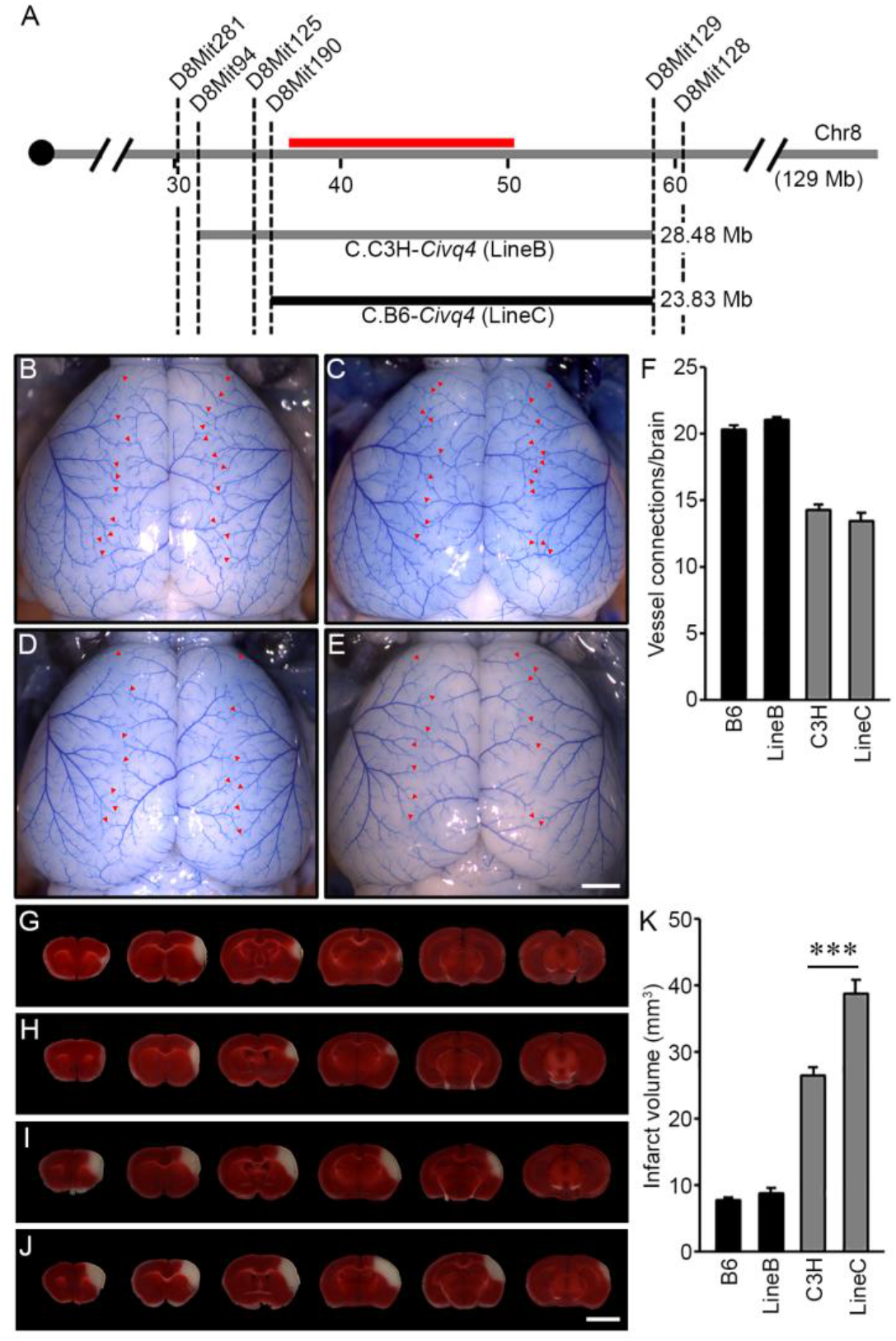
Validation of the location and allelic effects of the locus on chromosome 8 in congenic animals. **(A)** Schematic of the previously-mapped *Civq4* interval using reciprocal congenic mouse lines carrying either a segment of C3H chromosome 8 introgressed into the B6 background (C.C3H-*Civq4* (LineB)) or a segment of B6 chromosome 8 introgressed into the C3H background (C.B6-*Civq4* (LineC)). The red bar indicates the 1.5-LOD support interval of QTL mapping between B6 and C3H. Together, these reciprocal congenic mouse lines cover the entire interval of the *Civq4*. **(B – E)** Representative images of the brains for inbred mouse strains, B6 (B) and C3H (D), and reciprocal congenic mouse lines, LineB (C) and LineC (F). The red arrowheads indicate collateral vessel connections between the ACA and MCA. Scale bar: 1 mm. **(F)** The graph shows the average number of collateral vessel connections between the ACA and MCA in the brain. The total number of animals for B6, LineB, C3H, and LineC were 37, 30, 21, and 12 mice, respectively. Data represent the mean ± SEM. **(G – J)** Serial brain sections (1 mm) for both inbred strains, B6 (G) and C3H (I), and the reciprocal congenic mouse lines, LineB (H) and LineC (J), are shown. The infarct appears as white tissue after 2% TTC staining. Scale bar: 5 mm. **(K)** The graph indicates the infarct volume 24 hr after pMCAO. The total number of animals for B6, LineB, C3H, and LineC were 34, 35, 24, and 16 mice, respectively. Data represent the mean ± SEM. *** *p* < 0.001, 2-tailed Student’s *t* test.

Using two inbred mouse strains, B6 and C3H, and the reciprocal congenic mouse lines, LineB and LineC, we first analyzed the number of collateral vessel connections between the anterior cerebral artery (ACA) and middle cerebral artery (MCA). The number of vessel connections of the congenic mouse lines matched that of the parental lines for their overall strain background, and not the smaller congenic interval (B6 (20.4), LineB (21.0), C3H (14.3), LineC (13.5) (Fig 4B – F). Thus, as expected, the congenic regions did not affect the collateral vessel phenotype. We then measured the effects of the *Civq4* region on ischemic infarct volume. There was no difference in infarct volume between B6 and the LineB (introgressed C3H for the candidate region on the B6 background) (B6 (7.8 mm^3^) and LineB (8.5 mm^3^)). However, the infarct volume in LineC (introgressed B6 for the candidate region on the C3H background) displayed a significantly larger infarct volume (150%) after pMCAO compared to C3H (its background inbred mouse strain) (38.7 mm^3^ *vs*. 26.4 mm^3^) (Fig 4G – K). It is important to note that while there was no difference in collateral vessel number between the C3H and LineC mice (Fig 4F and K), infarct volume was increased in the LineC compared to its background C3H mouse strain. Therefore, the congenic lines confirm that a gene(s) mapping within the *Civq4* modulates infarct volume after cerebral ischemia through a collateral vessel-independent mechanism. Additionally, this confirms that the B6 allele in the *Civq4* acts as a risk allele for ischemic infarction.

### Coding SNP variation identifies potential candidate genes in the locus

To identify the causal gene(s) modulating infarct volume in the *Civq4* interval, we first surveyed all coding genes in the interval for the presence of non-synonymous coding SNPs (hereafter, coding SNP). Through this bioinformatic analysis, we identified a total of 964 coding SNPs in 62 coding genes within this interval. Indeed, only 10 genes mapping within the interval did not show a coding SNPs between the mapping strains. Thus, to further refine the candidate gene list, we filtered the gene list by an additional criterion, using the assumption at least one infarct modulating gene in the *Civq4* interval is also shared in the overlapping interval mapping between WSB and B6. We further required the allele present in the two protective strains (i.e., C3H and WSB allele) to differ from the risk strain (i.e., B6 allele). A total of 85 coding SNPs in 18 coding genes remained (S2 Table). Next, to determine whether an amino acid substitution within any of these coding SNPs has functional consequences on the respective protein, all the coding SNPs were evaluated using three independent *in silico* prediction algorithms, SIFT, PolyPhen-2, and PROVEAN. Through these *in silico* functional analyses, 8 genes within the *Civq4* were determined to have a coding SNP (or SNPs) predicted to be “damaging” by at least one of the prediction algorithms. Only one gene, *Zdhhc2*, harbors a coding SNP predicted to be damaging by all three algorithms and another two genes, *Prag1* and *Msr1*, harbor damaging coding SNPs predicted by two algorithms (Table 2).

**Table 2.**
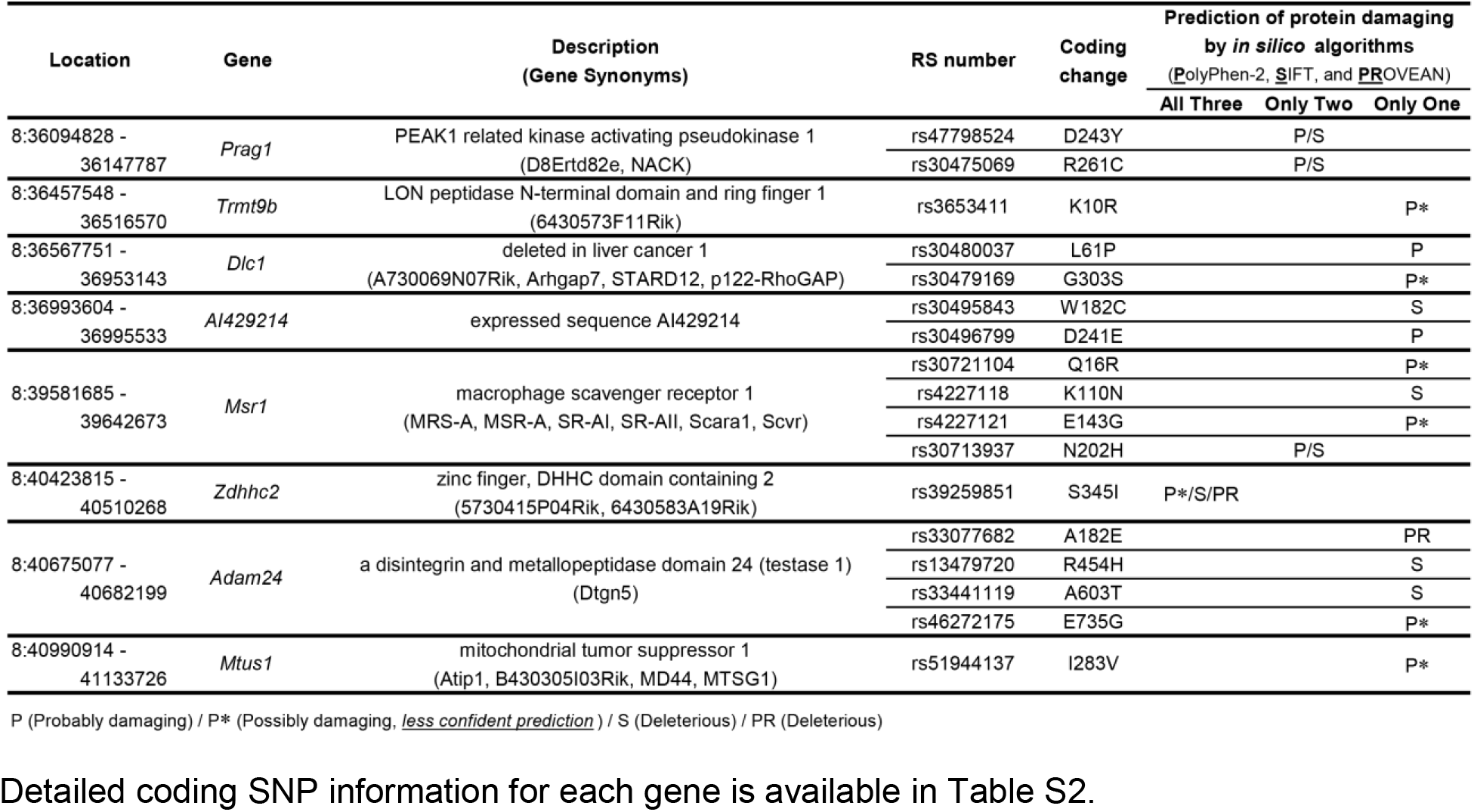
Candidate genes within the 1.5-LOD support interval harboring coding SNPs among the three inbred mouse strains.

### RNA Sequencing reveals additional candidate genes

In order to identify additional candidate genes within the locus, we compared strain-specific transcript levels from adult mouse brain cortex tissue between the strains used in these two independent crosses (WSB *vs*. B6 and C3H *vs*. B6). Differences could be caused by regulatory sequence variation acting in *cis* by any number of potential molecular mechanisms but would ultimately be observed as strain-specific differences in transcript levels. When comparing WSB and B6, a total of 1,716 genes showed statistically-significant different transcript levels for genes across the entire mouse genome, but only 34 of these are located within the 1.5-LOD support interval on chromosome 8 (Fig 5A and Table 3A). When comparing C3H and B6, a total of 2,180 genes showed statistically-significant different transcript levels in genes across the entire mouse genome, but only 10 of these are located within the *Civq4* (1.5-LOD support interval) on chromosome 8 (Fig 5B and Table 3B). Since the two genetic regions identified by independent crosses overlap on chromosome 8, we made an assumption that at least one potential infarct-modulating gene would be located within the overlapping region. Using this criterion, the candidate genes that are differentially expressed between WSB and B6 is reduced to only 8 genes (highlighted in gray in Table 3A). Among these genes only three genes (*Mtmr7, Gm5345*, and *Gm40493*) are differentially expressed between WSB *vs*. B6 and C3H *vs*. B6. *Gm5345* is annotated as a processed pseudogene and *Gm40493* is a novel, predicted gene without additional annotation. *Mtmr7* is the only of the three genes annotated as protein-coding. We note that all the genes showing different transcript levels within the *Civq4* interval are lower in both C3H and WSB compared to B6 (Table 3A and B). Details for all of the genes showing different transcript levels (1,716 for WSB *vs*. B6 or 2,180 for C3H *vs*. B6) are listed in S3 Table A and B, respectively.

**Table 3A.**
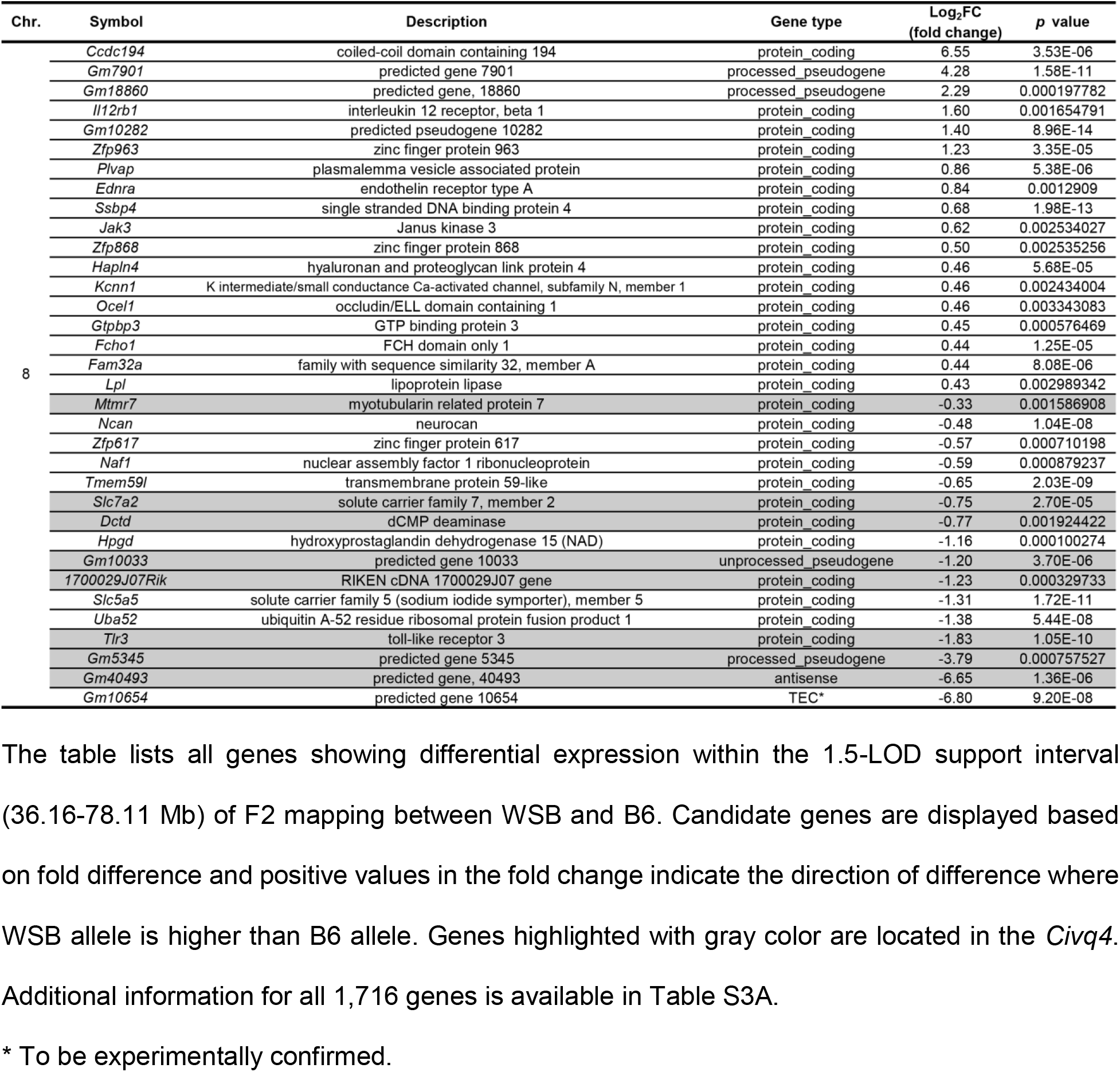
Differential gene expression between WSB and B6 determined by RNA sequencing analysis.

**Table 3B.** Differential gene expression between C3H and B6 determined by RNA sequencing analysis.

| Chr. | Symbol | Description | Gene type | Log<sub>2</sub>FC<br>(fold change) | p value |
| --- | --- | --- | --- | --- | --- |
| 8 | *Cnot7* | CCR4-NOT transcription complex, subunit 7 | protein_coding | -0.37 | 0.00227032 |
|  | *Mtmr7* | myotubularin related protein 7 | protein_coding | -0.38 | 0.000592403 |
|  | *Vps37a* | vacuolar protein sorting 37A | protein_coding | -0.42 | 0.002065146 |
|  | *Dlc1* | deleted in liver cancer 1 | protein_coding | -0.48 | 8.19E-05 |
|  | *Sorbs2* | sorbin and SH3 domain containing 2 | protein_coding | -0.50 | 0.000450996 |
|  | *Micu3* | mitochondrial calcium uptake family, member 3 | protein_coding | -0.60 | 0.000122213 |
|  | *Trmt9b* | tRNA methyltransferase 9B | protein_coding | -0.67 | 0.000345454 |
|  | *Sgcz* | sarcoglycan zeta | protein_coding | -1.06 | 0.003695057 |
|  | *Gm40493* | predicted gene, 40493 | antisense | -4.92 | 5.94E-05 |
|  | *Gm5345* | predicted gene 5345 | processed_pseudogene | -6.38 | 6.39E-06 |

**Fig 5.**
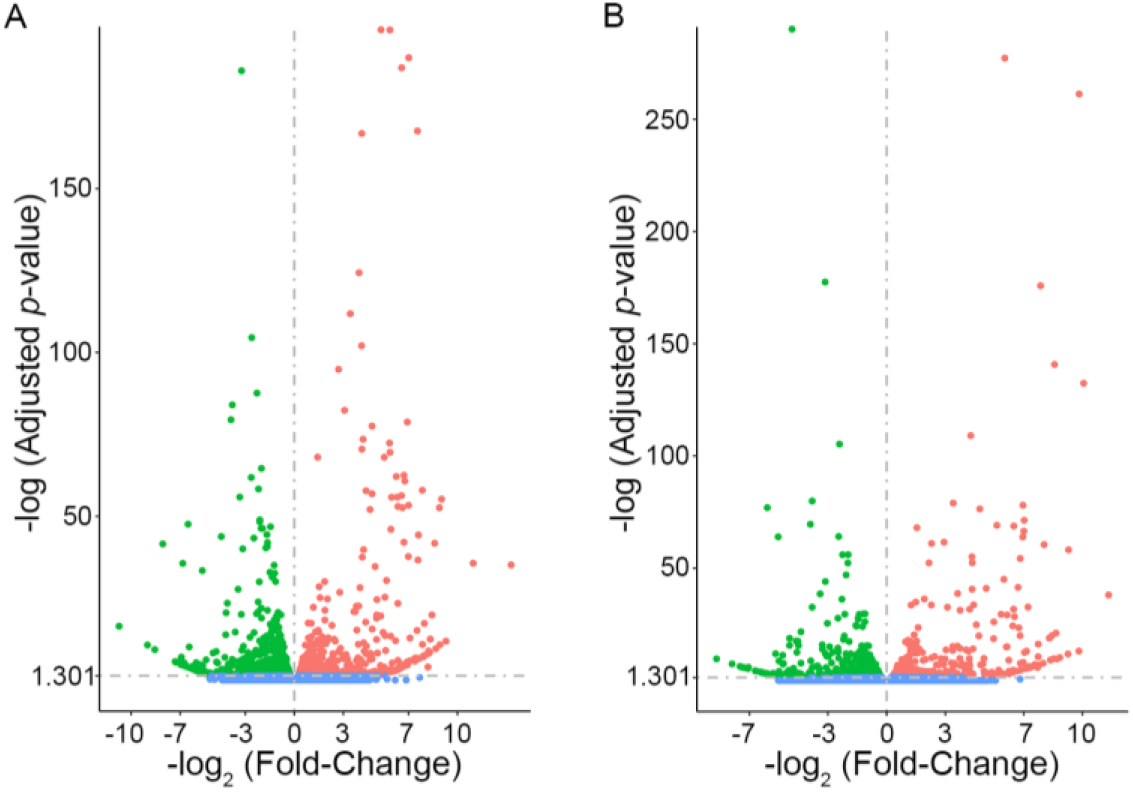
RNA sequencing identifies transcripts showing strain-specific differences in cerebral cortex gene expression. The volcano plots show differential gene expression for either WSB *vs*. B6 (A) or C3H *vs*. B6 (B) from brain cortex determined by RNA sequencing analysis. Each dot represents a different gene with log_2_ fold-changed plotted against log_10_ *p*-value. **(A)** The total number of significantly differentially expressed genes across the mouse genome between WSB and B6 is 1,716 genes, containing 782 upregulated genes indicated by red dots and 934 downregulated genes indicated by green dots, relative to B6. Blue dots (25,956) indicate no significant difference in genes compared between WSB and B6. **(B)** The total number of significantly differentially expressed genes across the mouse genome between C3H and B6 is 2,180 genes, containing 980 upregulated genes indicated by red dots and 1,200 downregulated genes indicated by green dots, relative to B6. Blue dots (25,433) indicate no significant difference in genes compared between C3H and B6.

### *Msr1* located in the *Civq4* plays a role in modulating ischemic stroke infarction

Through identifying genes either harboring coding SNPs predicted to be damaging (Table 2) or showing different transcript levels for both WSB *vs*. B6 and C3H *vs*. B6 (Table 3A and B), we uncovered potential candidate genes located in the *Civq4*. Of these candidate genes, *Msr1* shows multiple SNPs predicted to be damaging (Table 2 and S2 Table). In addition, recent studies report that *Msr1* plays an important role in cellular phagocytic activity in damaged tissue. In terms of the host defense mechanism, *Msr1* contributes to the pathology and repair of ischemic brain injury [23,24]. Therefore, we considered *Msr1* as a priority candidate gene for further study.

To investigate whether *Msr1* modulates ischemic infarct volume, we employed an *Msr1* KO mouse strain, representing a complete loss-of-function allele of this candidate gene. Although the *Msr1* KO mouse was originally generated in a mixed genetic background consisting of both 129×1/SvJ and C57BL/6J, the KO allele was backcrossed to B6 for over 12 generations and maintained at the Jackson Laboratory. Using whole genome SNP genotyping, we confirmed that this KO allele is fully contained in the B6 background. We then examined the effects of the loss of this gene on pial collateral vessel density. The number of vessel connections in the possible genotypes of *Msr1* KO allele mice did not differ among the genotypes (WT (21.1), Het (21.2), and KO (21.2)) (S6 Fig A), nor when compared to the parental background strain (B6 (20.4)) (Fig 1A). Thus, as expected, even complete loss of this gene has no effect on the collateral vasculature. We next measured ischemic infarct volume after pMCAO for each genotype of the *Msr1* KO mice. Infarct volume among genotypes also did not differ between WT (10.1 mm^3^), Het (10.3 mm^3^), and KO (8.4 mm^3^) (S6 Fig B), nor when compared to the parental background strain (B6 (7.8 mm^3^)) (Fig 1B). For a KO *Msr1* allele in the B6 background, this result was perhaps not too surprising. If the extensive collateral vessel connections in B6 enable rapid reperfusion of the ischemic territory, this could fully compensate for the lack of a neuroprotection due to loss of the *Msr1* gene. To test this hypothesis, we backcrossed the *Msr1* KO allele into the inbred strain C3H, which shows much fewer collateral vessel connections compared to that observed in the B6 strain (Fig 4F). We backcrossed the *Msr1* KO allele to N5 and validated the mice as nearly 100% C3H by whole genome SNP genotyping.

We then examined the effect of the loss of *Msr1* on pial collateral vessel density and infarct volume after pMCAO in the C3H background, which due to a paucity of collateral vessel connections, would reduce the reperfusion of the ishemic territory and thereby possibly blunt or even suppress the neuroprotective effect. As observed with the *Msr1* KO mice in the B6 background, the number of vessel connections in the genotypes of the *Msr1* KO allele on the C3H background showed no differences among the genotypes (WT (14.3), Het (14.2), and KO (12.9)) (Fig 6A – F and G). However, the infarct volume in the *Msr1* KO mice on the C3H background (37.7 mm^3^) was larger (157%) than infarct volume observed in WT mice (24.0 mm^3^) (Fig 6H, J, and K). The infarct volume in the genotypes of *Msr1* KO mice on the C3H background after pMCAO displayed a gene dosage effect with increased infarct volume (136%) in *Msr1* Het mice (32.7 mm^3^) compared to WT mice (Fig 6I and K). Moreover, the infarct volume after pMCAO in the *Msr1* KO mice (the loss of the gene effect) and congenic mouse LineC (a small genetic interval substitution) showed a similar level of increased infarct volume compared to WT littermates and C3H mice, respectively (Fig 4K and Fig 6K). Taken together, these data show that *Msr1* has an essential role in modulating ischemic stroke infarction independent of any effects of collateral circulation.

**Fig 6.**
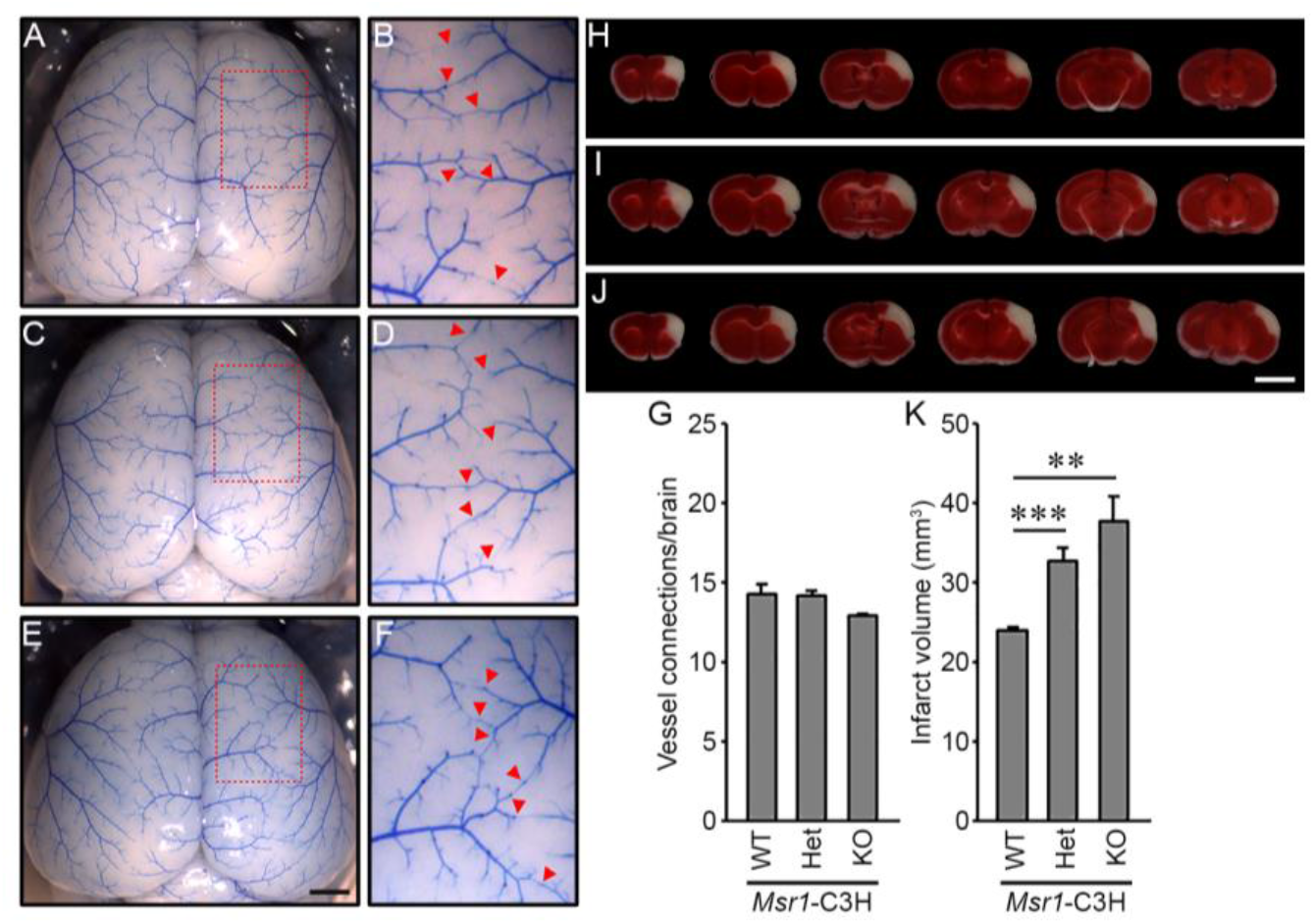
Collateral vessel anatomy and infarct volume after pMCAO in the *Msr1* KO mice. (A, C, and E) Representative images are shown for *Msr1*, wild-type (A), heterozygous KO (C), and homozygous KO (E) mouse strains in the C3H background. Scale bar: 1 mm. **(B, D, and F)** Each outlined area in a red box in A, C, and E is threefold magnified in B, D, and F, respectively, and red arrowheads indicate collateral vessel connections between the ACA and MCA. **(G)** The graph indicates the average number of collateral vessel connections in the brain. The total number of animals for *Msr1* WT, -Het, and –KO are 7, 14, and 10 animals, respectively. **(H – J)** Serial brain sections (1 mm) for each genotype of *Msr1* WT (H), Het (I), and KO (J) 24 hr after pMCAO. The infarct appears as white tissue after 2% TTC staining. Scale bar: 5 mm. **(K)** The graph shows the infarct volume for *Msr1* WT, -Het, and –KO. The total number of animals for *Msr1* WT, -Het, and –KO are 11, 20, and 16 animals, respectively. Data represent the mean ± SEM. ** *p* < 0.01 and *** *p* < 0.001, 2-tailed Student’s *t* test.

## Discussion

We have attempted to identify novel genes modulating infarct volume in ischemic stroke through a collateral vessel independent manner (i.e., via neuroprotection). Although the collateral circulation strongly affects brain tissue damage (infarction) after onset of stroke via reperfusion of the ischemic territory, we sought to identify strains that break the inverse correlation between collateral vessel density and infarct volume after stroke induction. We previously found one inbred strain, C3H, that exhibits a similar number of collateral vessel connections compared to B6, but these two strains show significantly different infarct volume after pMCAO [11]. Using F2 progeny between B6 and C3H, we performed genome-wide linkage analysis for infarct volume and uncovered a robust neuroprotective locus on chromosome 8 that was designated as *Civq4* [11]. Classical inbred mouse strains have been used extensively for QTL mapping of diverse phenotypic traits. These inbred mouse strains originally came from mixture of subspecies of the *Mus musculus* (*M. m*.), specifically; *M. m. musculus, M. m. domesticus*, and *M. m. castaneus* [25]. Unfortunately, the *M. m*. subspecies, *M. m. domesticus*, dominates the genetic background of the classical inbred mouse strains [26,27]. Thus, although these classical inbred mouse stains contain various pairwise combinations of their ancestral subspecies haplotypes, due to a predominance of genetic information from one subspecies they lack a full representation of genetic diversity for genetic mapping studies.

To expand the pool of genetic diversity among mouse strains, we previously added eight founder mouse strains of the Collaborative Cross [20] to the pool of our mapping strains. These genetically diverse founder strains include five classical inbred mouse strains (129S1/SvlmJ, A/J, B6, NOD/ShiLtJ, and NZO/HILtJ) and three wild-derived inbred mouse strains from different *M. m*. subspecies (CAST (*M. m. castaneus*), PWK/PhJ (*M. m. musculus*), and WSB (*M. m. domesticus*)) [28]. We identified one additional mouse strain (WSB) that exhibits a high number of collateral vessels, but demonstrates an increased infarct volume after pMCAO [20]. The WSB strain and another wild-derived strain (CAST) display the highest number of collateral vessel connections among all strains examined thus far. Therefore, we performed genome-wide QTL mapping analysis in an F2 progeny between these two wild-derived strains and discovered four novel neuroprotective loci that modulate ischemic stroke infarction in a collateral vessel-independent mechanism [20].

In this study, we attempted to discover additional neuroprotective loci by exploiting other strains that exhibit a disconnect between collateral vessel density and infarct volume. We crossed the commonly used classical inbred strain C57BL/6J, with the wild-derived strain, WSB. The infarct volume of the F2 progeny between these strains provides additional evidence of a collateral vessel-independent neuroprotective effect; while the number of collateral vessel connections are tightly distributed near their parental-strain phenotypes, infarct volumes vary across the entire phenotypic spectrum observed in other inbred mouse strains [10,20], (Fig 1A and B). In this F2 cross between B6 and WSB, we found a single statistically-significant locus mapping to chromosome 8. This locus overlaps a region on chromosome 8 *Civq4* that we previously identified from an F2 mapping between B6 and C3H. Both loci were identified with classical strain B6 crossed with another strain (C3H or WSB). Further evidence the loci may involve the same gene in the overlapping region on chromosome 8 is that for both loci, B6 is the susceptibility locus (Fig 3B and S5 Fig C), even when the parental strain overall is protective. Due to the time (>1 year) required to generate a fully congenic strain, we do not have validation data on reciprocal congenic lines between B6 and WSB for the newly mapped chromosome 8 locus. However, we do have data for the overlapping *Civq4* locus. Importantly, the *Civq4* reciprocal congenic mouse lines also support the direction of allele-specific phenotypic effects. LineC introgressed with a segment of the *Civq4* region from the B6 into the C3H background shows dramatically increased infarct volume after pMCAO compared to the C3H, whereas the LineB introgressed with a segment of the *Civq4* region from C3H into the B6 background exhibits a similar infarct volume compared to the B6. We surmise that MCAO creates a minimal region of cell death and that in B6 and Line B, this minimum has been reached. By contrast, collateral vessel connections in LineB and LineC show no difference compared to their background strains. Therefore, the overall evidence of two independent crosses suggests that a gene (or genes) on chromosome 8 regulates ischemic stroke infarction in a collateral vessel-independent manner. Assuming that the same gene is involved, we focused our attention on candidate genes located within the overlapping regions of the two loci. To include both protein-coding and potential regulatory variation, we compiled a list of candidate genes exhibiting either, strain-specific coding SNPs or transcript level differences. From the list of the candidate genes, we ranked *Msr1* as a high priority candidate gene based on *in silico* SNP analysis and biological plausibility. *Msr1* harbors 10 coding SNPs with differences between the protective and risk alleles, four of which are predicted to alter protein function. *Msr1* does not show strain-specific differential gene expression between either WSB *vs*. B6 or C3H *vs*. B6 comparisons. *Msr1* is mainly expressed in neonate microglia and other cells of the nervous system and microglial *Msr1* transcript is increased in Alzheimer’s disease [29,30]. More significant to this work, a recent study has shown that *Msr1* prevents the exacerbation of ischemic stroke damage by enhancing phagocytic activities in macrophage and microglia to resolve inflammation [24]. We therefore examined phenotypes of *Msr1* KO mice. As we expected for a collateral vessel-independent locus, there is no difference in the number of collateral vessel connections between the genotypes of *Msr1* KO and WT mice. By contrast, infarct volume after onset of stroke in *Msr1* KO mice is dramatically increased compared to WT. Furthermore, this infarct volume phenotype shows a gene dosage effect as the infarct volume of *Msr1* heterozygotes is larger than WT but smaller than KO mice. We note that the infarct volume phenotype of this gene is only observed in the *Msr1* KO mice on the C3H background. We reason that since the infarct volume in B6 strain is largely modulated by reperfusion due to robust collateral circulation seen in this strain, this effect compensates for the loss of *Msr1*. In other words, when ischemia is only transient due to reperfusion, the neuroprotective effect of other loci is blunted. While we examined *Msr1* based on our prioritization of candidate genes within the genetic interval on chromosome 8, further studies may identify additional genes mapping to this locus that modulate ischemic stroke infarction. Nonetheless, *Msr1* is one of the genes that modulates this effect.

In summary, by utilizing inbred mouse strain pairs that exhibit high collateral vessel connections but show large differences in infarct volume after stroke induction, we identified a genetic locus modulating ischemic stroke infarction in a collateral vessel-independent mechanism. This genetic region on chromosome 8 overlaps with the *Civq4* locus that we previously mapped in a previous cross. Using the reciprocal congenic mouse lines, we validated the phenotypic effects of this genetic interval on chromosome 8. *In silico* predictive coding SNP analysis and RNA sequencing data analysis provided a list of potential candidate genes. Using a knockout allele, we determine that *Msr1* influences infarct volume in a collateral independent manner. Further investigation may identify other genes that modulate infarct volume at this locus. The identification of these genes as modulators of infarct volume may provide potential therapeutic targets for human ischemic stroke.

## Materials and methods

### Animals

All inbred mouse strains and *Msr1* KO mice (B6.Cg-*Msr1*^*tm1Csk*^/J) were obtained from the Jackson Laboratory (Bar Harbor, ME), and then bred locally to obtain mice used in all experiments. To generate the reciprocal congenic mouse lines, F1 animals between B6 and C3H were backcrossed to either B6 for C.C3H-*Civq4* (LineB) or C3H for C.B6-*Civq4* (LineC). After confirmation of a segment of the *Civq4* region by genotyping, female reciprocal congenic mouse lines were backcrossed to the descendants ∼ 10 generations with male recipient stain. The *Msr1* KO mice in the B6 mouse background were backcrossed a minimum of five generations with C3H to generate the *Msr1* KO in the C3H strain background. The genetic background of both reciprocal congenic lines and *Msr1* KO mice were confirmed by whole genome SNP genotyping (OpenArray Technology, Waltham, MA). Mice (both male and female animals) were age-matched (P21 for collateral vessel density and 12 ± 1 week for pMCAO) for all experiments.

### Ethics statement

All animal study procedures were conducted with protocols approved by the Duke University IACUC (A105-16-05 and A081-19-04) in accordance with NIH guidelines.

### Collateral vessel density measurement

As we have shown that collateral vessel traits are established by 3 weeks of age and remain constant for many months [11,16], the collateral vessel phenotype was measured at P21 as previously described [12,13]. Mice were anesthetized with ketamine (100 mg/kg) and xylazine (2.5 mg/kg), and the ascending thoracic aorta was cannulated. The animals were perfused with freshly made buffer (1 mg/ml adenosine, 40 g/ml papaverine, and 25 mg/ml heparin in PBS) to remove the blood. The pial circulation was then exposed after removal of the dorsal calvarium and adherent dura mater. The cardiac left ventricle was cannulated and a polyurethane solution with a viscosity sufficient to minimize capillary transit (1:1 resin to 2-butanone, PU4ii, VasQtec) was slowly infused; cerebral circulation was visualized under a stereomicroscope during infusion. The brain surface was topically rinsed with 10% PBS-buffered formalin and the dye solidified for 20 min. After post-fixation with 10% PBS-buffered formalin, pial circulation was imaged. All collaterals interconnecting the anterior- and middle cerebral artery trees of both hemispheres were counted.

### Permanent MCAO

Focal cerebral ischemia was induced by direct permanent occlusion of the distal MCA as previously described [12,13]. Briefly, adult mice were anesthetized with ketamine (100 mg/kg) and xylazine (2.5 mg/kg). The right MCA was exposed by a 0.5 cm vertical skin incision midway between the right eye and ear under a dissecting microscope. After the temporalis muscle was split, a 2-mm burr hole was made with a high-speed micro drill at the junction of the zygomatic arch and the squamous bone through the outer surface of the semi-translucent skull. The MCA was clearly visible at the level of the inferior cerebral vein. The inner layer of the skull was removed with fine forceps, and the dura was opened with a 32-gauge needle. While visualizing under an operating microscope, the right MCA was electrocauterized. The cauterized MCA segment was then transected with microscissors to verify permanent occlusion. The surgical site was closed with 6-0 sterile nylon sutures, and 0.25% bupivacaine was applied. The temperature of each mouse was maintained at 37°C with a heating pad during the surgery until the animal was fully recovered from the anesthetic. Mice were then returned to their cages and allowed free access to food and water in an air-ventilated room with the ambient temperature set to 25°C.

### Infarct volume measurement

Cerebral infarct volumes were measured 24 h after surgery because the size of the cortical infarct is largest and stable at 24 h after distal permanent MCA occlusion [31]. Twenty-four hours after MCAO surgery, the animals were euthanized with CO_2_ inhalation followed by decapitation, and the brains were carefully removed. The brains were placed in a brain matrix and sliced into 1 mm coronal sections after being chilled at -80°C for 4 min to slightly harden the tissue. Each brain slice was placed in 1 well of a 24-well plate and incubated for 20 min in a solution of 2% 2,3,5-triphenyltetrazolium chloride (TTC) in PBS at 37°C in the dark. The sections were then washed once with PBS and fixed with 10% PBS-buffered formalin at 4°C. Then, 24 h after fixation, the caudal face of each section was scanned using a flatbed color scanner. The scanned images were used to determine infarct volume [32]. Image-Pro software (Media Cybernetics) was used to calculate the infarcted area of each slice by subtracting the infarcted area of the hemisphere from the non-infarcted area of the hemisphere to minimize error introduced by edema. The total infarct volume was calculated by summing the individual slices from each animal.

### SNP genotyping

Genomic DNA was isolated from tails of F2 progeny between WSB and B6 mice using DNeasy Tissue kit (Qiagen, Hilden, Germany). Genome-wide SNP genotyping was performed with a new Mouse Universal Genotyping Array (MiniMUGA), an array-based genetic QC platform with over 11,000 probes. Array hybridization including sample preparation was performed by Neogen/GeneSeek (Lincoln, NE).

### Quantitative Trait Locus (QTL) analysis

Using R/qtl software, genome-wide scans were performed as previously described [20] with minor changes. Genotype information from the MiniMUGA were prepared for QTL mappings as followed. A total of 515 informative markers for WSB and B6 across the mouse genome were used for genetic mapping. The significance thresholds for LOD scores were determined by 1,000 permutations using all informative markers. A QTL was considered significant when its LOD score exceeded 95% (*P* < 0.05) of the permutation distribution. The confidence interval of the peak was determined by the 1.5-LOD support interval. The physical map (megabase; Mb) positions based on the genomic sequence from the GRCm38/mm10 were calculated using Mouse Map Converter tool of the Jackson Laboratory (http://cgd.jax.org/mousemapconverter/). For the interval *Civq4*, SNP data were obtained from the Mouse Phenome Database (https://phenome.jax.org/).

### SNP analysis

SNPs between the strains that cause an amino acid change were predicted to be detrimental to a protein by three independents *in silico* prediction algorithms, Polymorphism Phenotyping v2 (PolyPhen-2, http://genetics.bwh.harvard.edu/pph2/index.shtml) [33], Sorting Intolerant From Tolerant (SIFT, http://sift.jcvi.org) [34], and Protein Variation Effect Analyzer (PROVEAN, http://provean.jcvi.org) [35].

### RNA sequencing analysis and differential gene expression

Paired-end, 150 bp sequencing reads were generated from adult (8 – 12 week) brain cortex tissue mRNA of B6, C3H, and WSB (3 male mice for each strain) mice on the NovaSeq 6000 instrument. Adapters for all paired-end sequencing reads were trimmed using Cutadapt [36] and gene expression counts for each sample were obtained using HTseq [37]. Differential gene expression analysis between two groups was performed using DESeq2 R package. DESeq2 provides statical routines for determining differential expression in digital gene expression data using a model based on the negative binominal distribution. *p*-values were adjusted using a false discovery rate of 5% by Benjamini-Hochberg correction [38]. These RNA sequencing and differential gene expression analyses were performed by Novogene (Sacramento, CA).

### Statistical analysis

Significant differences between data sets were determined according to the following definition: *p* > 0.05, not significant; *p* = 0.01 – 0.05, significant (*); *p* = 0.001 – 0.01, very significant (**); and *p* < 0.001, highly significant (***). All values were represented as the mean ± SEM.

## Acknowledgments

The authors thank Drs. Namsoo Kim and Sena Bae for helpful discussion concerning data sorting and analysis, and Mr. Christian R. Benavides for animal husbandary. This work was supported by a grant from NIH grant 5R01NS100866 and the Foundation Leducq Transatlantic Network of Excellence in Neurovascular Disease (17 CVD 03).

## Supporting information captions

**S1 Fig. Infarct volume in F2 animals shows relatively wide distribution**.

Each graph describes the distribution of infarct volume for B6, WSB, F1, F2 animals shown in Figure 1B with scatter plots. The range of the infarct volume is shown on the x-axis and the y-axis indicates animal numbers.

**S2 Fig. Non-parametric mapping test of the genome-wide QTL mapping analysis**.

The Non-parametric mapping test was performed to validate the genome-wide QTL mapping for infarct volume that was analyzed with the parametric mapping test shown in Figure 2. The non-parametric test indicated by a red line is superimposed on the parametric test indicated by a black line. The two independent mapping plots are nearly identical.

**S3 Fig. Characterization of the allelic effects of the locus mapping to chromosome 17**. The linkage peak on chromosome 17 did not reach a significance threshold (*p* < 0.05) determined by 1,000 permutation test. However, genotype-phenotype correlation of the F2 cohort at gUNC27989312 (LOD score: 3.51) in chromosome 17 shows that the B6 allele is protective for infarction. This follows the same trend as the parental strains. Data represent the mean ± SEM. *NS p* ≥ 0.05, ** *p* < 0.01, and *** *p* < 0.001, 2-tailed Student’s *t* test.

**S4 Fig. Infarct volume after pMCAO in the three inbred mouse strains used in two independent QTL mapping studies that identified an overlapping interval on chromosome 8**.

The same data of infarct volume for each inbred strain are shown in either Fig 1B (for B6 and WSB) or Fig 4K (for B6 and C3H). The graph shows the infarct volume 24 h after pMCAO. The total number of animals for B6, C3H, and WSB are 34, 24, and 18 animals, respectively. Data represent the mean ± SEM.

**S5 Fig. Allelic effects of the Civq4 on chromosome 8 mapping previously identified in a cross between C3H and B6**.

**(A)** The graph presents the re-analysis of a genome-wide QTL mapping scan for infarct volume measured 24 h after pMCAO using 210 F2 animals (B6 and C3H). Chromosomes 1 through X are represented numerically on the x-axis and the y-axis represents the LOD score. The significant (*p* < 0.05) level of linkage was determined by 1,000 permutation tests. Three regions of the genome mapping to chromosome 5, 8, and 18 display significant QTL mapping to the infarct volume trait with LOD scores of 4.13, 10.31, and 4.71, respectively. **(B)** The graph shows the major locus mapping across chromosome 8 using 14 informative SNP markers. The LOD score at the peak is 10.31 (rs13479735), and the 1.5-LOD support interval is from 36.02 to 50.26 Mb, indicated by the red bar on the graph. **(C)** Genotype-phenotype correlation of the F2 cohort at rs13479735. The B6 allele confers increased susceptibility to infarction and the C3H allele confers protection. Data represent the mean ± SEM. ** *p* < 0.01 and *** *p* < 0.001, 2-tailed Student’s *t* test.

**S6 Fig. Collateral vessel anatomy and infarct volume after pMCAO in the *Msr1* KO mice on the B6 background**.

**(A)** The graph indicates the average number of collateral vessel connections in the brain. The total number of animals for *Msr1* WT, -Het, and –KO are 14, 28, and 18 animals, respectively. Data represent the mean ± SEM. **(B)** The graph shows the infarct volume for *Msr1* WT, -Het, and –KO. The total number of animals for *Msr1* WT, -Het, and –KO are 19, 27, and 22 animals, respectively. Data represent the mean ± SEM.

**S1 Table. Genotype and phenotype information of 376 F2 (B6 x WSB) animals**.

The information was used for genome-wide QTL mapping analysis. (XLSX)

**S2 Table. Genes mapping within the *Civq4* that harbor coding SNP differences between protective allele and risk allele**.

The table shows non-synonymous coding SNPs between the protective allele (i.e., C3H and WSB) and the risk allele (i.e., B6). The functional consequences on protein function for each coding SNP was predicted using three independent *in silico* algorithms, SIFT, PolyPhen-2, and PROVEAN. Coding SNP predicted to be either “damaging” or “deleterious” are highlighted in red. (XLSX)

**S3 Table A. Genes show ing strain-specific differential gene expression between WSB and B6**.

For each of the 1,716 genes, the table displays the *p*-value and fold change, with either a positive value or negative value for increased or decreased expression of the WSB allele, relative to the B6 allele. (XLSX)

**S3 Table B. Genes show strain-specific differential gene expression between C3H and B6**. For each of the 2,180 genes, the table displays the *p*-value and fold change, with either a positive value or negative value for increased or decreased expression of the C3H allele, relative to the B6 allele. (XLSX)

## References

1. Bogousslavsky J, Aarli J, Kimura J. Stroke: time for a global campaign? Cerebrovasc Dis. 2003;16:111–3.

2. WHO: Global burden of stroke (https://www.who.int/cardiovascular_diseases/en/cvd_atlas_15_burden_stroke.pdf).

3. Johnson W, Onuma O, Owolabi M, Sachdev S. Stroke: a global response is needed. Bull. World Health Organ. 2016;94(9):634.

4. Benjamin EJ, Muntner P, Alonso A, Bittencourt MS, Callaway CW, Carson AP, et al. Heart Disease and Stroke Statistics—2019 Update: A Report From the American Heart Association. 2019;139(10):e56–e528.

5. Roger VL, Go AS, Lloyd-Jones DM, Benjamin EJ, Berry JD, Borden WB, et al. Heart disease and stroke statistics--2012 update: a report from the American Heart Association. Circulation. 2012;125(1):e2–e220.

6. Jamrozik K, Broadhurst RJ, Anderson CS, Stewart-Wynne EG. The role of lifestyle factors in the etiology of stroke. A population-based case-control study in Perth, Western Australia. Stroke. 1994;25(1):51–9.

7. Dichgans M. Genetics of ischaemic stroke. Lancet Neurol. 2007;6(2):149–61.

8. Malik R, Chauhan G, Traylor M, Sargurupremraj M, Okada Y, Mishra A, et al. Multiancestry genome-wide association study of 520,000 subjects identifies 32 loci associated with stroke and stroke subtypes. Nat Genetics. 2018;50:524–37.

9. Keum S, Marchuk DA. A locus mapping to mouse chromosome 7 determines infarct volume in a mouse model of ischemic stroke. Circ Cardiovasc Genet. 2009;2(6):591–8.

10. Keum S, Lee HK, Chu PL, Kan MJ, Huang MN, Gallione CJ, et al. Natural genetic variation of integrin alpha L (Itgal) modulates ischemic brain injury in stroke. PLoS Genet. 2013;9(10):e1003807.

11. Chu PL, Keum S, Marchuk DA. A novel genetic locus modulates infarct volume independently of the extent of collateral circulation. Physiol Genomics. 2013;45(17):751–63.

12. Lee HK, Keum S, Sheng H, Warner DS, Lo DC, Marchuk DA. Natural allelic variation of the IL-21 receptor modulates ischemic stroke infarct volume. J Clin Invest. 2016;126(8):2827–38.

13. Lee HK, Koh S, Lo DC, Marchuk DA. Neuronal IL-4Rα modulates neuronal apoptosis and cell viability during the acute phases of cerebral ischemia. FEBS J. 2018;285(15):2785–98.

14. Wang S, Zhang H, Wiltshire T, Sealock R, Faber JE. Genetic Dissection of the Canq1 Locus Governing Variation in Extent of the Collateral Circulation. PLoS One. 2012;7(3):e31910.

15. Lucitti JL, Sealock R, Buckley BK, Zhang H, Xiao L, Dudley AC, et al. Variants of Rab GTPase-Effector Binding Protein-2 Cause Variation in the Collateral Circulation and Severity of Stroke. Stroke. 2016;47(12):3022–31.

16. Chalothorn D, Clayton JA, Zhang H, Pomp D, Faber JE. Collateral density, remodeling, and VEGF-A expression differ widely between mouse strains. Physiol Genomic. 2007;30:179–91.

17. The Complex Trait Consortium. The Collaborative Cross, a community resource for the genetic analysis of complex traits. Nat Genet. 2004;36(11):1133–7.

18. Threadgill DW. Meeting report for the 4th annual Complex Trait Consortium meeting: from QTLs to systems genetics. Mamm. Genome. 2006;17(1):2–4.

19. Aylor DL, Valdar W, Foulds-Mathes W, Buus RJ, Verdugo RA, Baric RS, et al. Genetic analysis of complex traits in the emerging Collaborative Cross. Genome Res. 2011;21(8):1213–22.

20. Lee HK, Widmayer SJ, Huang MN, Aylor DL, Marchuk DA. Novel neuroprotective loci modulating ischemic stroke volume in wild-derived inbred mouse strains. Genetics. 2019;213(3):1079–92.

21. Kelada SNP, Aylor DL, Peck BCE, Ryan JF, Tavarez U, Buus RJ, et al. Genetic analysis of hematological parameters in incipient lines of the collaborative cross. G3. 2012;2(2):157–65.

22. Widmayer SJ, Handel MA, Aylor DL. Age and Genetic Background Modify Hybrid Male Sterility in House Mice. Genetics. 2020;216(2):585–97.

23. Kelley JL, Ozment TR, Li C, Schweitzer JB, Williams DL. Scavenger receptor-A (CD204): A two-edged sword in health and disease. Crit Rev Immunol. 2014;34(3):241–61.

24. Shichita T, Ito M, Morita R, Komai K, Noguchi Y, Ooboshi H, et al. MAFB prevents excess inflammation after ischemic stroke by accelerating clearance of damage signals through MSR1. Nat Med. 2017;23(6):723–32.

25. Wade CM, Daly MJ. Genetic variation in laboratory mice. Nat. Genet. 2005;37:1175–80.

26. Frazer KA, Eskin E, Kang HM, Bogue MA, Hinds DA, Beilharz EJ, et al. A sequence-based variation map of 8.27 million SNPs in inbred mouse strains. Nature. 2007;448(30):1050–3.

27. Yang H, Wang JR, Didion JP, Buus RJ, Bell TA, Welsh CE, et al. Subspecific origin and haplotype diversity in the laboratory mouse. Nat Genet. 2011;43(7):648–55.

28. Collaborative Cross Consortium. The genome architecture of the Collaborative Cross mouse genetic reference population. Genetics. 2012;190(2):389–401.

29. Husemann J, Loike JD, Anankov R, Febbraio M, Silverstein SC. Scavenger receptors in neurobiology and neuropathology: their role on microglia and other cells of the nervous system. Glia. 2002;40:195–205.

30. Husemann J, Silverstein SC. Expression of scavenger receptor class B, type I, by astrocytes and vascular smooth muscle cells in normal adult mouse and human brain and in Alzheimer’s disease brain. Am J Pathol. 2001;158:825–32.

31. Lambertsen KL, Meldgaard M, Ladeby R, Finsen B. A quantitative study of microglial-macrophage synthesis of tumor necrosis factor during acute and late focal cerebral ischemia in mice. J Cereb Blood Flow Metab. 2005;25(1):119–35.

32. Wexler EJ, Peters EE, Gonzales A, Gonzales ML, Slee AM, Kerr JS. An objective procedure for ischemic area evaluation of the stroke intraluminal thread model in the mouse and rat. J Neurosci Methods. 2002;113(1):51–8.

33. Adzhubei IA, Schmidt S, Peshkin L, Ramensky VE, Gerasimova A, Bork P, et al. A method and server for predicting damaging missense mutations. Nat Methods. 2010;7(4):248–9.

34. Ng PC, Henikoff S. SIFT: predicting amino acid changes that affect protein function. Nucleic Acids Res. 2003;31(13):3812–4.

35. Choi Y, Sims GE, Murphy S, Miller JR, Chan AP. Predicting the Functional Effect of Amino Acid Substitutions and Indels. PLoS One. 2012;7(10):e46688.

36. Martin M. Cutadapt removes adapter sequences from high-throughput sequencing reads. EMBnet J. 2011;17:10–2.

37. Anders S, Pyl PT, Huber W. HTSeq–a Python framework to work with high-throughput sequencing data. Bioinformatics. 2015;31:166–9.

38. Benjamini Y, Hochberg Y. Controlling the false discovery rate: a practical and powerful approach to multiple testing. J. R. Stat. Soc. B. 1995;57:289–300.

